# Multiple stage hydra effect in a stage–structured prey–predator model

**DOI:** 10.1101/2021.07.16.452738

**Authors:** Michel Iskin da S. Costa, Lucas dos Anjos, Pedro V. Esteves

## Abstract

In this work, we show by means of numerical bifurcation that two alternative stable states exhibit a hydra effect in a continuous–time stage–structured predator–prey model. We denote this behavior as a stage multiple hydra effect. This concomitant effect can have significant implications in population dynamics as well as in population management.

## Introduction

Positive effects at the population level from increasing mortality in consumer populations are sometimes called hydra effect (Abrams, 2009). Its definition is the increase in the equilibrium or time–averaged density of a consumer population with increasing mortality. This positive effect of mortality can have a strong appeal in applied ecology, for example, because harvest/removal of species is an additional form of mortality impinged on individuals (Abrams, 2009).

In a previous work (Costa and Anjos, 2018) we showed by numerical bifurcation that in the presence of bistability in a continuous–time predator–prey model with Allee effect in the predator, the two alternative stable states exhibited a concomitant hydra effect in the predator population. We then coined this behavior as multiple hydra effect.

The novelty in this short communication with respect to Costa and Anjos (2018) is that this multiple hydra effect is now shown to occur in the prey adult population of a stage–structured prey– predator model. Therefore, we denote this behavior as a stage multiple hydra effect. The appearance of this phenomenon in two distinct predator-prey community modules (Holt, 1997) indicates that it might be worthwhile to investigate whether the multiple hydra effect is a relatively common phenomenon in theoretical multispecies dynamical models of other community modules.

### A stage–structured prey–predator dynamical model

In this work, the biological setup we analyze consists of a stage–structured prey–predator where young and adult prey individuals are connected via maturation and fecundity. It is assumed that two types of harvest/predation act simultaneously on the adult prey population. This biological framework is shown schematically in figure 1 (hereinafter *FR*_*i*_ denotes functional response type *i* (*i* = *2, 3*) throughout this work).

**Figure 1:**
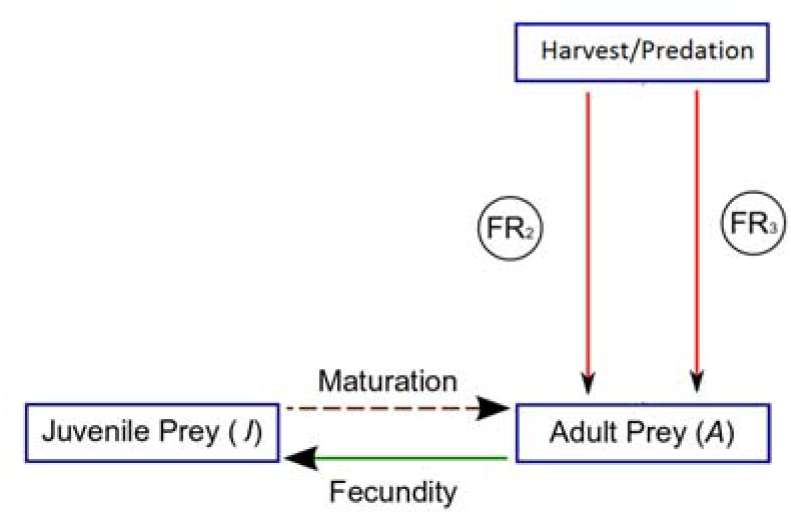
A scheme of a stage–structured prey–predator population where harvest/predation concomitantly acts upon the adult population with *FR*_*2*_ and *FR*_*3*_. Vertical arrows denote harvest/predation.

A continuous–time dynamical model based on Costa et al. (2017) for the trophic scheme of figure 1 can be given by:

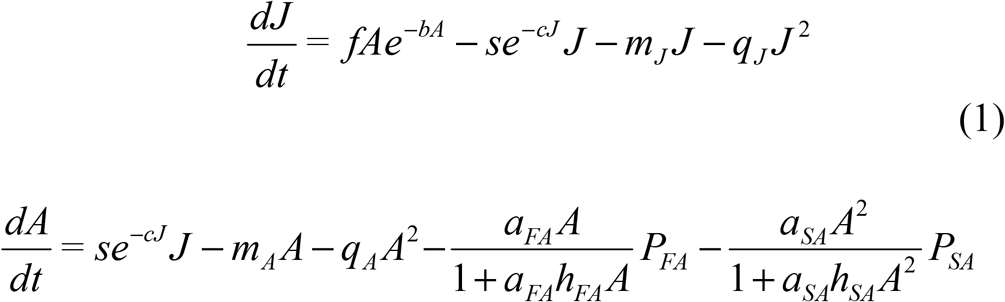

*J* is the density of prey young individuals and *A* is the density of prey adult individuals. The definition of variables and parameters of the model (1) are displayed in Table 1.

**Table 1.** Definition of variables and parameters of the model (1)

|  | Meaning |
| --- | --- |
| **Variables** |  |
| $J$ | young prey |
| $A$ | adult prey |
| **Parameters** |  |
| $f$ | maximum fecundity of adults |
| $b$ | density dependence in fecundity |
| $c$ | density dependence in maturation rate |
| $s$ | maximum value of the *per capita* maturation rate |
| $m_J$ | density-independent *per capita* mortality rate of young |
| $m_A$ | density-independent *per capita* mortality rate of adults |
| $q_J$ | coefficient of density-dependent *per capita* mortality rate of young |
| $q_A$ | coefficient of density-dependent *per capita* mortality rate of adults |
| $a_{FA}$ | harvest/predation attack coefficient on adults |
| $h_{FA}$ | manipulation time of harvest/predation |
| $P_{FA}$ | density/numbers of predator/harvest engines |
| $a_{SA}$ | harvest/predation attack coefficient on adults |
| $h_{SA}$ | manipulation time of harvest/predation |
| $P_{SA}$ | density/numbers of predator/harvest engines |

The model (1) assumes that competitive interactions occur within each prey stage (adults only compete with adults and juveniles with juveniles). The assumed density dependence in fecundity and maturation in the model (1) is based on the fact that the two density–dependent functions are plausible for many populations, where adult density dependence is often expressed in a stock–recruitment relationship, and juvenile competition is likely to reduce the growth rate to the adult stage (Abrams and Quince, 2005). Moreover, the reason for considering the combination of density–dependent functions in fecundity and maturation is its ability to produce alternative equilibrium for the prey population in the absence of the harvest/predation (i.e., *P*_*FA*_ = *P*_*SA*_ = 0 (Abrams and Quince, 2005)),. This feature is likely to complicate the overall dynamics when adding the harvest/predation disturbance (i.e., when *P*_*FA*_, *P*_*SA*_ > 0).

In terms of harvest/predation, *a*_*FA*_ *A*/(1+*a*_*FA*_ *T*_*hFA*_ *A*) is a type 2 functional response while *a*_*SA*_ *A*^*2*^/(1+*a*_*SA*_ *T*_*hSA*_ *A*^*2*^) is a type 3 functional response. These two expressions could well describe a commercial and a recreational fishery, respectively. Functional response type 2 can be appropriate to describe a commercial fishery because it exerts a strong fishing pressure when the fish density is low. On the other hand, functional response type 3 can be appropriate to describe a recreational fishery because it exerts a very weak fishing pressure when the target fish density is low. One reason for this behavior is that anglers are prone to turn to other types of fish when their target fish density is low. This theoretical fishery setup is in line with the suggestions to analyze harvest strategies in fisheries with functional responses types 2 and 3 (nonlinear functional responses) besides functional responses type 1 (a linear response sometimes named proportional harvest in fisheries) (Mangel (2005); but see also Beddington and Cooke (1982)).

Increased population size caused by increased mortality is known as a hydra effect (Abrams, 2009). In general, the analyses are based upon the variation of the *per capita* density–independent mortality rate of the species in case. However, it is important to stress that in our work the analysis of the hydra effects hinges upon the variation of the intensity of a density–dependent *per capita* mortality rate. Particularly, an increase in the prey adult population size (*A*) as the result of its increased mortality rate induced by the harvest/predation term *P*_*SA*_ *a*_*SA*_ *A*^*2*^/(1+*a*_*SA*_ *T*_*hSA*_ *A*^*2*^) can be called a stage hydra effect in the prey adult population of the model (1). Hence, to investigate whether this effect occurs in the model (1), we analyze the influence of *P*_*SA*_ on the prey adult population sizes in the model (1). Note that *P*_*SA*_ can be seen as the intensity of the density dependent *per capita* mortality rate *P*_*SA*_ *a*_*SA*_ *A*/(1+*a*_*SA*_ *T*_*hSA*_ *A*^*2*^) induced on adult prey by the functional response type 3. A similar procedure was employed in the stage–structured model analyzed in Costa et al. (2017).

The following analysis consists of one–parameter bifurcation diagram of the proposed model (1), which is drawn by means of the software package XPPAUT (Ermentrout, 2002). This software calculates the equilibrium points of the nonlinear differential equations system (1) together with the real part of their corresponding eigenvalues. This information is gathered to build the graphs displayed throughout this work. The resulting bifurcation diagram of the adult population in the model (1) as a function of *P*_*SA*_ is displayed in figure 2.

**Figure 2:**
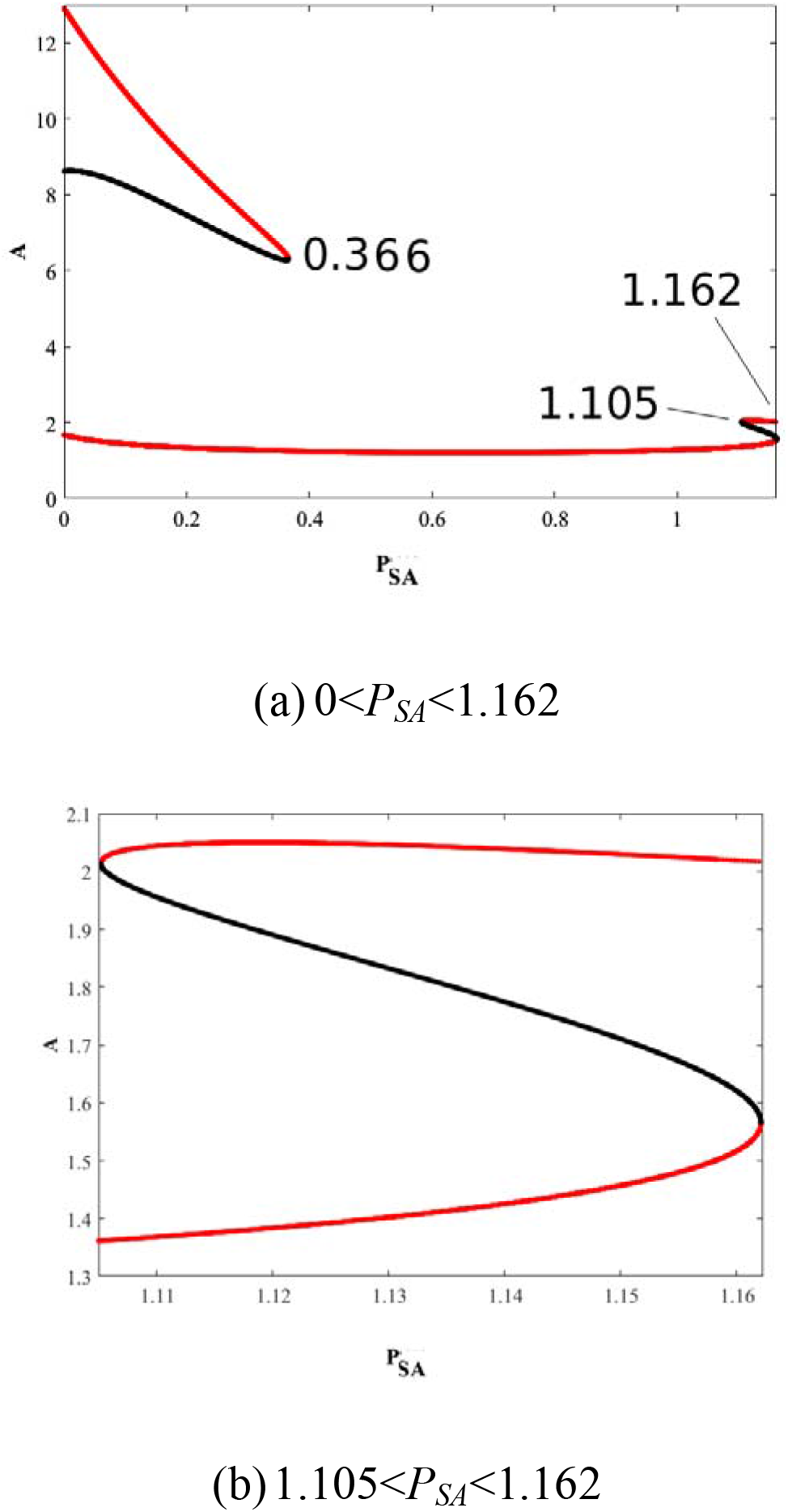
Bifurcation diagram of adults (*A*) in model (1) as a function of *P*_*SA*_. (a): the complete bifurcation diagram; (b) a zoom of the multiple hydra effect region. Red lines: stable equilibrium points; black lines: unstable equilibrium points. Parameter values: *f =* 0.35; *b =* 0.265; *s = 1; m*_*J*_ *=* 0.04; *q*_*J*_ *=* 0.005; *m*_*A*_ *=* 0.0008; *c=* 1; *q*_*A*_ = 0.00035; *a*_*FA*_=0.2; *h*_*FA*_=1; *P*_*FA*_=*0*.*1*; *a*_*SA*_=0.1; *h*_*SA*_=1.

Figure 2 shows that there are two locally stable equilibrium points for 0 < *P*_*SA*_ *<*0.37. For *P*_*SA*_ =0.37 there is a collapse bringing about the disappearance of the equilibrium point of the upper branch. For 1.105 < *P*_*SA*_ *<*1.1621 there appear again two locally stable equilibrium points, and they both increase for 1.105 < *P*_*SA*_ *<*1.122 – a phenomenon coined multiple hydra effect (Costa and Anjos, 2018). For *P*_*SA*_ >1.1621 there is a new collapse bringing about the disappearance of one locally equilibrium point (the one lying on the lower branch). It is important to remark that in predator–prey theory the lower stable branch is usually named predator pit and is commonly found when predation of prey by a constant population of predator is conveyed by a functional response type 3 (Turchin (2003); for the appearance of this phenomenon in a similar model to (1) see Costa et al (2017)).

As mentioned before, in applied ecology the multiple hydra effect in figure 2 can have significant implications in population management. In fisheries, for instance, it says that the magnitude of the fishery yield will depend on the initial stages of the fish population. This, in turn, demands a fairly accurate stock assessment when one aims for profitable economic returns from harvest. From the theoretical ecology standpoint, it is argued that exploring dynamics of an age–structured prey population model consumed by generalist, age–specific predators with a constant density over time is essential to help develop predator–prey theory (Pavlova and Berec, 2012). We think that the previous results in Costa and Anjos (2018) and the ones found in our present work contribute in this direction by showing multiple hydra effect in specific non-structured and stage–structured prey–predator models consisting of system of nonlinear differential equations.

## Discussion

The main result of this work concerns the appearance of a multiple hydra effect in the prey adult population in a structured prey–predator model described by a system of nonlinear coupled differential equations. In other words, the increase of the coefficient of density–dependent *per capita* mortality rate of the predator (*P*_*SA*_) along a specific interval generates bistability, and the two branches of this bistability show an increase in the adult prey population, giving rise to a stage multiple hydra effect. To our knowledge, we are not aware of such result in stage–structured population models consisting of nonlinear systems of autonomous differential equations. However, it is important to mention that several hydra effects (including the so–called multiple hydra effect presented in this work) are discussed for a seasonal non-structured population model described by a nonlinear system of difference equations in Liz (2017).

It is important to stress that in our work the hydra effect occurred by virtue of the variation of a density–dependent *per capita* mortality rate and not a density–independent *per capita* one because we intended to assess the effects of *P*_*SA*_ on the dynamics of a stage– structured–prey population. Hence, we resorted to a broader definition which relates the hydra effect to the increase of a species density with the increase of its mortality rate (Schröder et al. (2014)). Note that each stage of the prey population in the model (1) has three sources of mortality. In the context of predator–prey theory, for instance, we chose *P*_*SA*_ *a*_*SA*_ *A*^*2*^/(1+*a*_*SA*_ *T*_*hSA*_ *A*^*2*^) in order to assess how generalists predators (*P*_*SA*_ and *P*_*FA*_) with their functional responses influence the dynamics of a stage–structured prey population.

The occurrence of multiple hydra effect in instances of prey-predator models may point to the investigation of such behavior in other community module models (Holt, 1997) such as shared predator, omnivory, in order to assess its possible ubiquity in more complex food webs. This endeavor would certainly help contribute to the development of the food web population dynamics theory.

## References

Abrams, P. A., Quince, C. 2005. The impact of mortality on predator population size and stability in systems with stage– structured prey. Theoretical Population Biology, 68, 253–266.

Abrams, P. A. 2009. When does greater mortality increase population size? The long history and diverse mechanisms underlying the hydra effect. Ecology Letters, 12, 462–474.

Beddington, J. R., Cooke, J. G., 1982. Harvesting from a predator–prey complex. Ecological Modelling 14, 155–177.

Costa M.I.S., Esteves, P. V., Faria, L. D. B., Anjos, L. 2017. Prey dynamics under generalist predator culling in stage structured models. Mathematical Biosciences, 285, 68–74.

Costa, M. I. S., Anjos, L. 2018. Multiple hydra effect in a predator–prey model with Allee effect and mutual interference in the predator. Ecological Modelling, 373, 22–24.

Ermentrout, B. 2002. Simulating, analyzing, and animating dynamical systems: a guide to XPPAUT for researchers and students, Vol. 14, Philadelphia, PA, USA.

Holt, R. D. 1997. Community modules. In: Gange, A. C., Brown, V. K. (Eds.), Multitrophic interactions in terrestrial systems. Blackwell Science Oxford, UK, pp. 333–339.

Liz, E., 2017. Effects of strength and timing of harvest on seasonal population models: stability switches and catastrophic shifts. Theoretical Ecology, 10, 235–244.

Mangel, M. 2006. The theoretical biologist’s toolbox: quantitative methods for ecology and evolutionary biology. Cambridge University Press, Cambridge.

Pavlova, V., Berec, L. 2012. Impacts of predation on dynamics of age–structured prey: Allee effects and multi–stability. Theoretical Ecology, 5, 533–544.

Schröder, A., van Leeuwen, A., Cameron, T.C. 2014. When less is more: positive population–level effects of mortality. Trends in Ecology and Evolution, 29,11, 614–624.

